# Tactile attention modulates occipital alpha oscillations in early blindness

**DOI:** 10.1101/2022.06.03.494660

**Authors:** Ane Gurtubay-Antolin, Ricardo Bruña, Olivier Collignon, Antoni Rodríguez-Fornells

## Abstract

Alpha oscillatory activity is thought to contribute to the cueing of visual attention through the engagement of task-relevant occipital regions. In early blindness, occipital alpha oscillations are systematically reduced, suggesting that occipital alpha depends on visual experience. However, it is still unknown if alpha activity could serve attentional cueing in non-visual modalities in blind people, especially considering previous research that showed the recruitment of the occipital cortex for non-visual processing. To test this idea, we used electroencephalography to answer whether alpha oscillations reflected a differential recruitment of task-relevant regions between expected and unexpected conditions in two (texture and shape discrimination) haptic tasks. As expected, time frequency analyses showed that alpha suppression in parieto-occipital sites was significantly reduced in early blind individuals. The source reconstruction analysis revealed that group differences originated in the middle occipital cortex. In that region, expected trials evoked higher alpha desynchronization than unexpected trials in the EB group only. Our results support the role of alpha rhythms in the recruitment of occipital areas also in early blind participants, and for the first time we show that even if posterior alpha activity is reduced in blindness, it is however sensitive to task-dependent factors. Our findings therefore suggest that occipital alpha activity may be involved in tactile attention in blind individuals, maintaining the function proposed for visual attention in sighted population but switched to the tactile modality. Altogether, our results indicate that attention-dependent modulation of alpha oscillatory activity does not depend on visual experience.

**SIGNIFICANCE STATEMENT:** Are posterior alpha oscillations and their role in attention dependent on visual experience? Our results show that tactile attention can modulate posterior alpha activity in blind (but not sighted) individuals through the engagement of occipital regions, suggesting that in early blindness, alpha oscillations maintain their proposed role in visual attention but subserve tactile processing. Our findings bring a new understanding to the role that alpha oscillatory activity plays in blindness, contrasting with the view supporting that alpha activity is rather task unspecific.

## INTRODUCTION

Since the first electroencephalographic recordings, a close relationship has been suggested between visual processes and posterior (parieto-occipital) alpha oscillatory activity (Berger, 1929). Even if at first alpha oscillatory activity was considered to reflect cortical idling (Adrian & Matthews, 1934; Pfurtscheller et al., 1996), alpha oscillations are now thought to play an important role shaping the state of sensory regions directing the flow of information and allocating resources (Jensen & Mazaheri, 2010). Specifically, alpha activity is thought to contribute to attention (Foxe et al., 1998; Fu et al., 2001; Pfurtscheller & Lopes da Silva, 1999) enhancing the neural signal-to-noise ratio by reducing (i.e., desynchronizing) low frequency fluctuations (Hanslmayr et al., 2016; Mitchell et al., 2009) through the engagement of task-relevant neural populations (Jensen et al., 2012; Klimesch, 2012). In this line, alpha rhythms have been extensively related to visual attention. Specifically, they have been proposed to index visuospatial attention bias (Thut, 2006) and reflect anticipatory states of visual attention (Foxe et al., 1998; Rihs et al., 2009) as well as distractor suppression (Kelly et al., 2006; Rihs et al., 2007, 2009). Likewise, their lateralization has been found to vary with spatial attention (Kizuk & Mathewson, 2017; Rihs et al., 2007; Thut, 2006) and decreased alpha power during periods of expected target appearance has been related to improved performance (Min et al., 2008; Rohenkohl & Nobre, 2011).

However, it is still unknown to which extent posterior alpha oscillations and their role in gating attention depend on visual experience. A unique way to answer whether attention-induced changes could also modulate alpha oscillations regardless visual experience is to investigate posterior alpha oscillations in blind people. In early blind population, it has been consistently shown that posterior occipito-parietal regions reorganize to engage in the processing of non-visual inputs, both in the tactile (Amedi et al., 2003; Jiang et al., 2015; Sadato et al., 1998, 2004; van Kemenade et al., 2014) and auditory (Collignon et al., 2011; Dormal et al., 2016; Mattioni et al., 2020; Poirier et al., 2006; Rezk et al., 2020) modalities. If alpha activity is tightly linked to vision, it should therefore be altered in congenitally blind individuals. Several studies have actually suggested that this might be the case by showing that in early blindness, posterior alpha oscillations are decreased while participants are at rest (Adrian & Matthews, 1934; Hawellek et al., 2013; Noebels et al., 1978; Novikova, 1973), sleeping (Bertolo et al., 2003) or during haptic tasks (Kriegseis et al., 2006; Schepers et al., 2012; Schubert et al., 2015). The variety of tasks in which EB participants showed these reductions led to suggest that alpha rhythms did not play a role in sensory/cognitive functions, in the reorganized occipital cortex of blind people (Kriegseis et al., 2006). Consequently, no study so far directly tested whether attentional demands (e.g., trials matching or mismatching an expectancy) can modulate posterior alpha activity in the blind, a key signature of alpha oscillations in sighted people for visual processing (Foxe et al., 1998; Kizuk & Mathewson, 2017; Rihs et al., 2007, 2009; Thut, 2006).

In the present study, we used high-density EEG to answer whether different attentional conditions could modulate alpha oscillations in visually deprived people, by comparing alpha band activity in expected and unexpected conditions in two (texture and shape discrimination) haptic tasks. Given the prominent role of posterior alpha oscillations in the gating of visual attention through the recruitment of task-relevant regions (in the sighted) and considering that i) in early blind people occipito-parietal regions reorganize to engage in the processing of tactile stimuli (Beauchamp et al., 2007; Blake et al., 2004; Jiang et al., 2015; van Kemenade et al., 2014) and that 2) that occipital regions maintain their computational specialization following the loss of vision but shift their preferred input modality (Dormal & Collignon, 2011), we expected to find attention-induced alpha modulations in the early blind group.

## MATERIALS AND METHODS

The description of the sample of participants and the experimental procedure were previously reported in detail (Gurtubay-Antolin & Rodríguez-Fornells, 2017). Here, we reiterate the elements that are essential for understanding the results of the present study and describe in more detail those methods that are specific to the currently presented analyses.

### Participants

14 congenitally blind (7 women, mean ±SD, age = 35.7 ± 10.9 years) and 15 sighted participants (9 women, mean age = 29.3 ± 9.0 years) took part in the experiment. Both groups were matched by age and years of education. The inclusion criteria for the EB group included right handedness, less than 3% of visual residual abilities, blindness onset before 5 years of age, no recollection of visual memories and the ability to avoid blinks and control eye movements for 3 seconds. Blindness of cerebral origin was an exclusion criterion. Three congenitally blind participants were excluded from the analysis. EB4 was removed due to excessive muscular artifacts. EB10 was rejected due to residual abilities to read with a very high contrast and magnifiers, despite reporting 3% of residual visual abilities. EB14 was removed because he did not perform the shape discrimination task for timing reasons. The experiment was undertaken with the understanding and written consent of each participant and was approved by the local ethics committee in accord with the declaration of Helsinki. Demographic characteristics of early blind participants and control samples can be found in (Gurtubay-Antolin & Rodríguez-Fornells, 2017).

### Experimental procedure

The experimental procedure is reported in detail elsewhere (Gurtubay-Antolin & Rodríguez-Fornells, 2017). Briefly, participants conducted haptic shape and texture discrimination tasks with the right arm. In the texture-task, ten textures were used (cotton, cork, sackcloth, sandpaper, sponge, scourer, corduroy, suede, paper and velvet). In the shape-task, ten 2D wooden geometrical shapes were used (racket, circle, square, triangle, arrow, flower, crown, heart, star and lightning). Sighted participants were not allowed to see the objects during the entire experimental study, since we obstructed vision by placing an opaque screen between the subject and the item.

The texture- and shape-discrimination tasks had the same procedure (for illustration, see Figure 1 in Gurtubay-Antolin et al., 2017). To minimize individual differences in the haptic exploration, all participants conducted the same constrained exploration which consisted of contacting the object with three fingertips for ∼3 sec. Different constrained explorations were used between the tasks. In the texture-experiment, all three fingers were tied together and participants touched the textures with the fingertips, after descending the fingers through a vertical stick. In the shape discrimination task, participants touched the shapes at three specific locations (*contact points*), after sliding the fingers (which moved independently) through three rails that were carved into the table.

**Figure 1.**
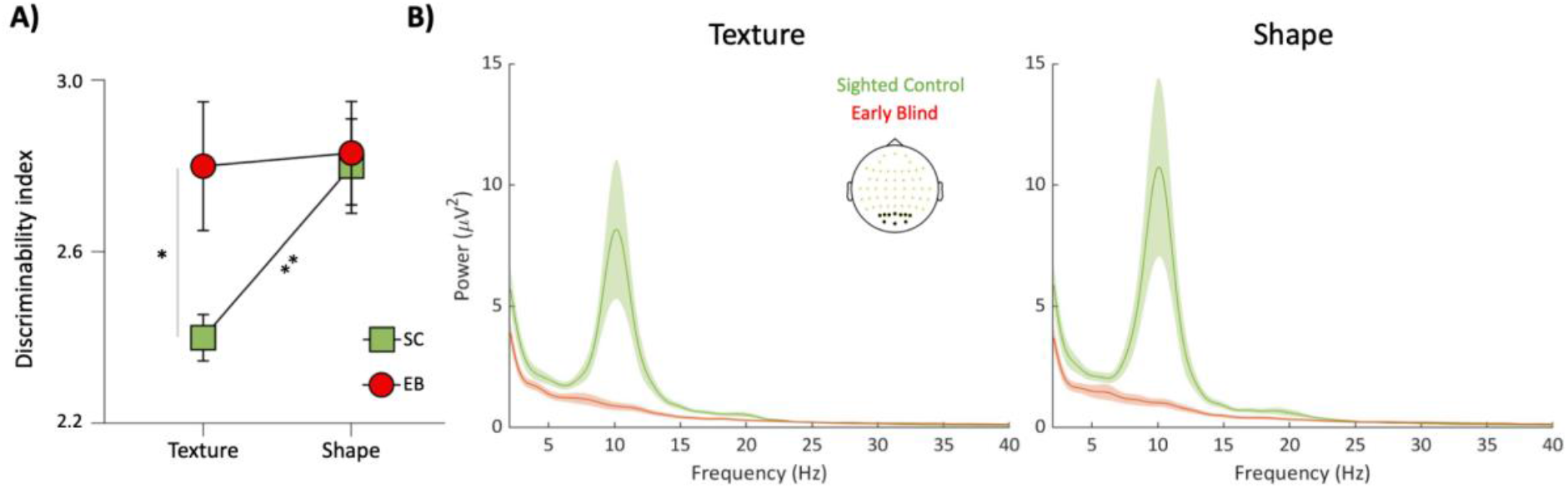
A. Differences in the discriminability index depending on the task and group. Notice the superior discriminability index of the EB group in the texture discrimination task. SC: Sighted Control group; EB: Early blind group. **p < 0.005, *p < 0.0125. Adapted from Gurtubay-Antolin et al., 2017. B. Average power spectral density in the posterior electrodes (marked in the electrode plot) for each group in the texture (left) and shape (right) discrimination tasks. The solid line represents the group average, while the shadowed line represents the standard error of the mean. The EB group shows reduced alpha activity in in posterior sites in both discrimination tasks.

Each trial began with the fingers in the initial position. Then, participants heard the name of an expected stimulus through the headphones. An auditory cue (a beep) indicated that the fingers could start moving towards the figure. In half of the trials the presented stimuli matched the previous expectancy (congruent condition), and in the other half it did not (incongruent). A second auditory cue indicated that a response had to be emitted, requesting the participants to press (with the left hand) a different keyboard button if the stimulus that they were touching corresponded to the expectancy or if not. Participants were requested to delay 3 s their motor response in order to avoid contamination of the EEG signal from motor related ERP-components.

The experimental session consisted of 180 trials conducted in 4 different blocks, interleaved by resting periods. Each block consisted of three series of 15 trials. To measure the time at which subjects contacted the object (contact time) the position and velocity of the fingers was recorded using an infrared motion capture system (CMS-30P, Zebris, Isny, Germany). The spatial resolution of the motion tracking system was 0.1 mm and a sampling frequency of 200 and 66 Hz was used in the texture and shape discrimination task, respectively. The difference in the sampling frequency between tasks comes from the fact that the three finger moved like an entity in the texture experiment (hence a single sensor was needed), whereas they moved independently in the shape-task (therefore three sensors were needed).

The instant in time when the participant touched the object was labeled as Contact-time (CT). CT was defined as the time when the absolute velocity of the first finger to contact the object was lower than 5% of its peak velocity. In the texture discrimination task, the arrival of the three fingers to the stimulus happened at the same time. On the contrary, the three fingers moved independently in the shape discrimination task. No further movement was allowed after contact-time, isolating the somatosensory processing in the ERPs from the previous motor execution. Trials with an incorrect Yes-No response (incorrect trials) or with a response-time higher than 2 s were removed from the analysis.

### Behavioral analysis

As described in (Gurtubay-Antolin & Rodríguez-Fornells, 2017), we conducted a two-way repeated measures ANOVA on the discriminability index with within-subject factor ‘group’ (Sighted Controls vs. Early Blinds) and between-subject variable ‘task’ (texture vs. shape).

### EEG acquisition

EEG recordings were acquired from 64 scalp electrodes (Electro-Cap International) using Brain-Vision Recorder software (version 1.3; Brain Products, Munich, Germany). Electrode positions were based on the standard 10/20 positions: Fpz/1/2, AF3/4, Fz/1/2/3/4/5/6/7/8, Fcz/1/2/3/4/5/6, Cz,/1/2/3/4/5/6, T7/8, Cpz/1/2/3/4/5/6, Tp7/8, Pz/1/2/3/4/5/6/7/8, POz/1/2/3/4/5/6, Oz/1/2. An electrode placed at the lateral outer canthus of the right eye served as an online reference. EEG was re-referenced offline to the average of the reference electrodes (Offner, 1950). Electrode impedances were kept below 5 kΩ. The EEG signal was sampled at 250 Hz and filtered with a bandpass of 0.03–45 Hz (half-amplitude cut-offs). Trials with base-to-peak electrooculogram (EOG) amplitude of more than 100 μV, amplifier saturation, or a baseline shift exceeding 200 μV/s were automatically rejected. These criteria removed electrocardiogram (ECG) contamination in most of the participants. After visually inspecting all the trials and in order to remove trials with remaining ocular artifacts, independent component analysis (ICA) (Delorme et al., 2007) was conducted in three subjects where the previous rejection was not enough to remove ECG *contamination*.

### Time-frequency analysis

Data was analyzed using MATLAB (The Mathworks, Inc., Munich, Germany), EEGLAB (Delorme & Makeig, 2004) and FieldTrip (Oostenveld et al., 2011). The epoch was defined from 2 seconds prior to the contact-time to 2.5 after it, and time-frequency (TF) activity was evaluated using a 7-cycle complex Morlet function. Following previous reports on reduced alpha waves in blind population (Adrian & Matthews, 1934; Noebels et al., 1978; Novikova, 1973), we first conducted a sanity check where we measured overall alpha activity on occipital sites during the baseline (−500 to 0 ms). We expected to replicate previous results on decreases of the posterior alpha activity in the EB group. We then assessed task-effects on alpha band activity in epochs between −0.2 and 1.0 s, in steps of 20 ms, and between 1 and 40 Hz, in linear steps of 1 Hz. Trials of the same condition [congruent, incongruent and all (congruent + incongruent)] were averaged for each subject and baseline corrected using the relative change before performing a grand average with all the individuals. The baseline was defined as 200 ms prior to the contact-time (−200 to 0 ms).

### Source reconstruction analysis

Since Time-Frequency analyses are not informative of the location where possible group differences arise, we performed a source reconstruction analysis using a template head model. We parted from the New York head (Huang et al., 2016, p. 201), extracted the brain, skull, and scalp masks, and created a three-layer boundary elements model with their interfaces. Last, we combined this model with the electrode positions provided by the manufacturer of the EEG system and created a lead field using OpenMEEG (Gramfort et al., 2010). As source model we used an homogeneous volumetric grid, with 1 cm of spacing between sources, and labeled the sources according to the automated anatomical labeling (AAL) atlas (Tzourio-Mazoyer et al., 2002). Only the 1210 source positions labeled as cortical were used.

As an inverse method we used dynamic imaging of coherent sources (DICS) beamformer (Gross et al., 2001), and adaptive spatial filter based on the cross-spectral density (CSD) matrix. To maximize the quality of the reconstruction, we tuned the filter using the average CSD matrix for the frequencies and latencies where we found differences in the TF analysis. We performed the source reconstruction separately for each participant and condition, using a participant-specific spatial filter built from the “all” condition (common filter approach). Last, we reconstructed the average power at each source position for the frequencies and latencies of interest, and corrected by the average power in the same source position during baseline, using the relative change.

### Statistical analysis

We performed statistical analyses in three levels, each feeding from the information obtained in the previous one. Firstly, we compared the time-frequency activity between groups (Sighted Control vs. Early Blind) and between conditions (Congruent vs. Incongruent) using a nonparametric cluster-based analysis (Maris & Oostenveld, 2007), defining clusters from the frequential, temporal, and spatial adjacency. The result of this first analysis is a range of frequencies and latencies where the activity between groups or conditions differs. To assess the location where group differences arose, we then reconstructed the source-level activity for the significant results, this is, limiting our reconstruction to these frequencies and latencies, and repeated the nonparametric analysis in source space, defining clusters from the spatial adjacency. The result of this analysis is a spatial set of adjacent source positions showing differences in those frequencies and latencies. Lastly, we computed post-hoc analyses to assess the effect of the task in each group in the source space. We defined a region of interest (ROI) from these sources and introduced its average activity into a mixed-effects ANOVA model with one between-subjects factor (the group), one within-subjects factor (the condition) and their interaction.

### Nonparametric cluster-based statistics

The nonparametric statistics used in the first and second levels of analysis in this work are based on the selection of clusters of adjacent elements (this is, frequencies, latencies, or positions) with a consistent behavior not explainable by chance. The procedure is as follows: 1) a statistical contrast, in this case either a one sample or two samples t-test, is performed for each data bin (frequency, latency, and position); 2) the significant results (p < 0.01) are clustered by adjacency, and the resulting cluster receives a value (the cluster statistic) equal to the sum of the t-statistic of all its members; 3) the data is resampled into random partitions, and 1 and 2 are repeated, generating a null distribution of cluster statistics; and 4) the statistic obtained in the original cluster is compared to this null distribution, and a *p*-value is assigned to it accordingly. In this work we used 10,000 permutations for the sensor-level statistics and 100,000 permutations for the source-level statistics.

## RESULTS

### Behavioral results

The behavioral results were previously reported in (Gurtubay-Antolin & Rodríguez-Fornells, 2017). Here, we reiterate the findings that are essential for understanding the results of the present study. We found a main effect of task with a higher discriminability index in the shape discrimination task compared to the texture discrimination task (F(1,25) = 6.31, p = 0.019, η^2^_p_ = 0.20) and a significant interaction (F(1,25) = 4.86, p = 0.037, η^2^ _p_ = 0.16). Post-hoc t-tests revealed significant differences in the control group (t(14) = 3.51, p = 0.003, Cohen’s d = 1.26) with a lower discrimination index for textures (d’ = 2.4 ± 0.2) than for shapes (d’ = 2.8 ± 0.4). No differences between tasks were observed in the EB group (t(11) = 0.01, p = 0.99, Cohen’s d = 0). In addition, independent-sample t-tests between groups revealed a higher discriminability index in the texture task for EBs when compared to controls (t(25) = 2.75, p = 0.011, Cohen’s d = 1.05, EBs d’ = 2.8 ± 0.5; Controls d’ = 2.4 ± 0.2) and no differences were found between groups for the shape discrimination task (t(25) = 0.06, p = 0.95, Cohen’s d = 0) (see Figure 1A).

### Time frequency analysis

As a sanity check, we first replicated previous reports on reduced alpha waves in blind population (Adrian & Matthews, 1934; Noebels et al., 1978; Novikova, 1973). We calculated the total alpha power (between 8 and 12 Hz) during the −500 −0 ms interval in occipital and parieto-occipital electrodes and we compared it between groups. The results showed that Early Blind individuals have significantly lower alpha power during both texture (t(25) = 2.76, p = 0.0109, Cohen’s d = 1.18) and shape discrimination (t(25) = 2.70, p = 0.0124, Cohen’s d = 1.16) tasks that the Sighted Control group (see Figure 1B).

We then assessed task-effects on alpha band activity in epochs between −0.2 and 1.0 s and between 1 and 40 Hz. From here on, all the power estimates will be relative to the baseline (−200ms, 0). In the texture discrimination task, the comparison between groups (Sighted Control vs. Early Blind) showed a significant cluster (p = 0.0274) in the alpha band (10 −12 Hz) between 260 and 560 ms after the contact, where the Sighted Control group showed lower alpha power compared to the Early Blind group (see Figure 2A). In these frequencies and latencies we found that both groups presented a desynchronization relative to baseline (for texture discrimination task, EB group t(10) = −2.30, p = 0.0443, Cohen’s d = −0.69; Controls t(14) = −8.00, p < 0.0001, Cohen’s d = −2.07; for shape discrimination task, EB group t(10) = −3.88, p = 0.0030, Cohen’s d = −1.17; Controls t(14) = −11.51, p < 0.0001, Cohen’s d = −2.97), with the control group showing significantly larger values of desynchronization. The effect appeared in occipital, left temporal, and frontal electrodes. The comparison between conditions (Congruent vs. Incongruent) showed no statistically significant differences at this level.

**Figure 2.**
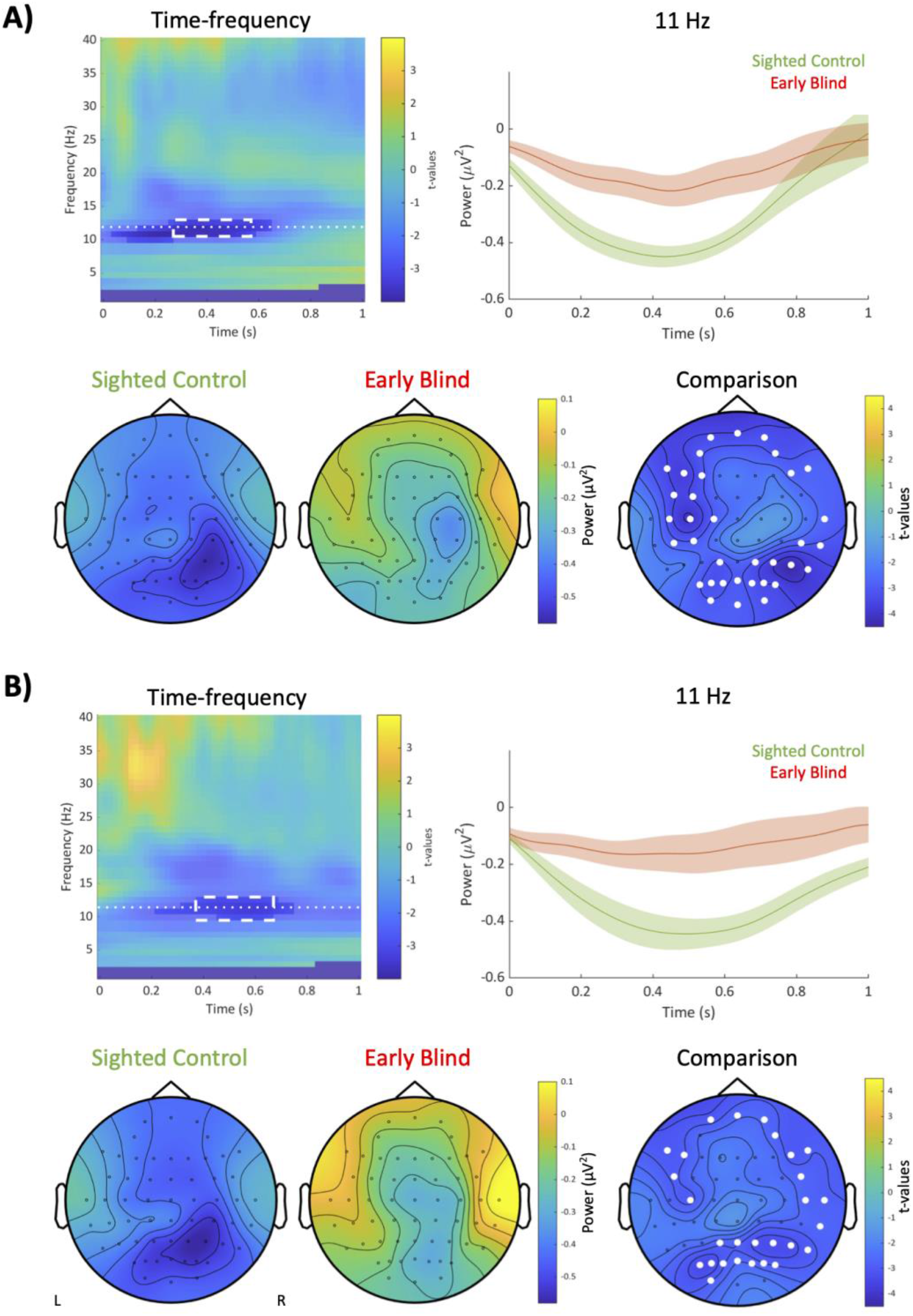
Results of the time-frequency analyses for the texture (A) and shape (B) discrimination tasks. The top-left graph shows the average time frequency over all electrodes, with the significant cluster marked in opaque colors and the nonsignificant data in semi-transparent colors. The top-right graph shows the evoked-related synchronization/desynchronization, separately for the Sighted Control (green) and the Early Blind (red) groups for the wavelet centered at 11 Hz (center of the significant data, represented by a dotted line in the time-frequency graph). The bottom-left graph shows the topographic representation of the power for the frequencies and times of interest (represented by a dashed rectangle in the time-frequency graph), separately for the Sighted Control and the Early Blind groups. The bottom-right graph shows the topographic representation of the differences between groups (t-statistic) for the frequencies and times of interest. Notice the reduced alpha desynchronization in the Early Blind group -compared to the Sighted Control- and the suppression relative to baseline in both groups (in both tasks).

In the shape discrimination task we found similar results. The comparison between groups (Sighted Control vs. Early Blind) showed a significant cluster (p = 0.0284) in the alpha band (9 - 12 Hz) between 360 and 660 ms after the contact, where again the Sighted Control group showed lower alpha power (see Figure 2B). Both groups showed a desynchronization respect to baseline, with the Sighted Control group presenting significantly higher values of desynchronization. Again, the effect appeared in occipital, temporal, and frontal areas. We did not find any significant result in the comparison between conditions.

### Source reconstruction analysis

To investigate where the effects seen in the Time-Frequency analysis originate from (their location), we reconstructed in source space the average power between 10 and 12 Hz in the latencies between 260 and 560 ms for the texture discrimination task, and between 9 and 12 Hz in the latencies between 360 and 660 ms for the texture discrimination task. We then calculated the between-groups comparisons in the source space. As we expected a lower power relative to baseline in the Sighted Control group, we performed a one-tailed t-test.

In the texture discrimination task, the results showed a significant cluster (p = 0.0156) where the Sighted Control individuals showed a higher desynchronization than the Early Blind individuals. The comparison between both groups in the shape discrimination task showed similar results, with one significant cluster (p = 0.0174). As depicted in Figure 3, in both cases the largest differences between groups were located in occipital areas, especially the bilateral middle occipital lobe, according to the AAL atlas. Therefore, we selected these areas for the post-hoc analyses.

**Figure 3.**
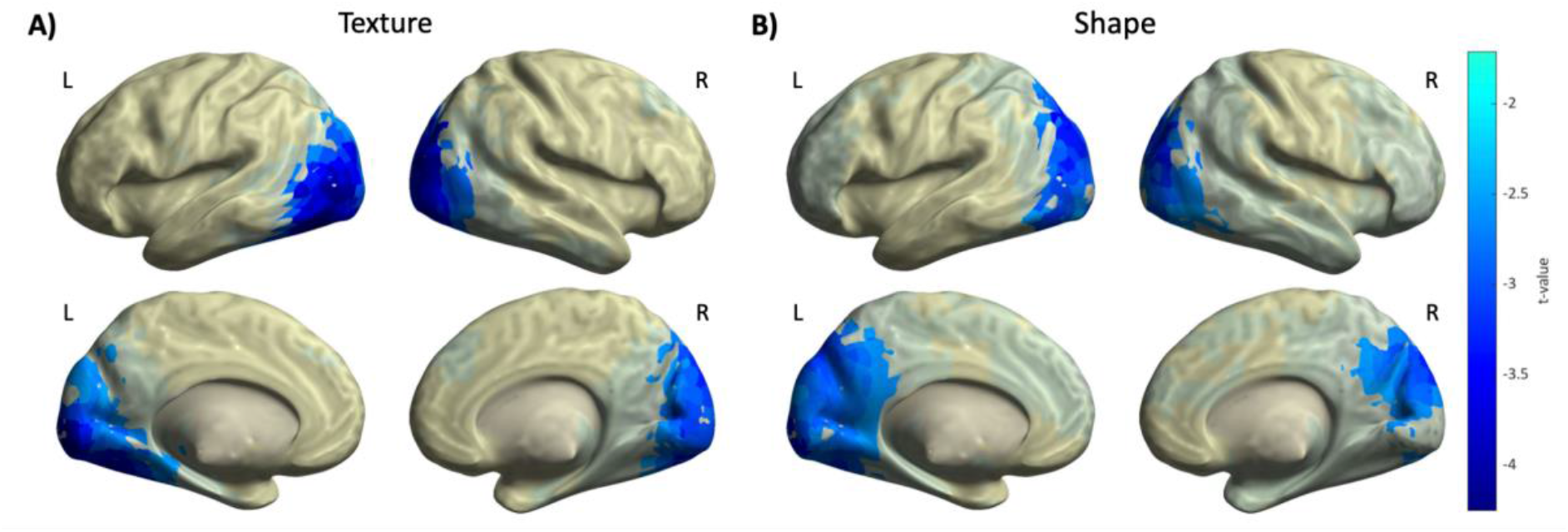
Topographic representation of the differences between groups (t-statistic) for the frequencies and times of interest in source space for the texture (A) and shape (B) discrimination tasks. Largest differences between groups were located in occipital areas, specially in the bilateral middle occipital lobe. L: Left, R: Right.

### Post-hoc analyses

We computed post-hoc analyses to assess the effect of the task in each group in the source space. As previously indicated, the post-hoc analyses consisted on an ANOVA contrast where the average power in the middle occipital lobe, relative to baseline, is compared between groups and conditions.

When evaluating the texture discrimination task, the ANOVA contrast revealed both a significant effect of group (F_1,24_ = 12.9, p = 0.0015) and condition (F_1,24_ = 12.3, p = 0.0018), together with a marginally significant interaction between both factors (F_1,24_ = 4.23, p = 0.0509). When considering both groups separately, the effect of condition appeared in the Early Blind individuals (p = 0.0007), where the Congruent condition showed significantly higher desynchronization values than the Incongruent one, but not in the Sighted Control individuals. These results are shown in Figure 4A.

**Figure 4.**
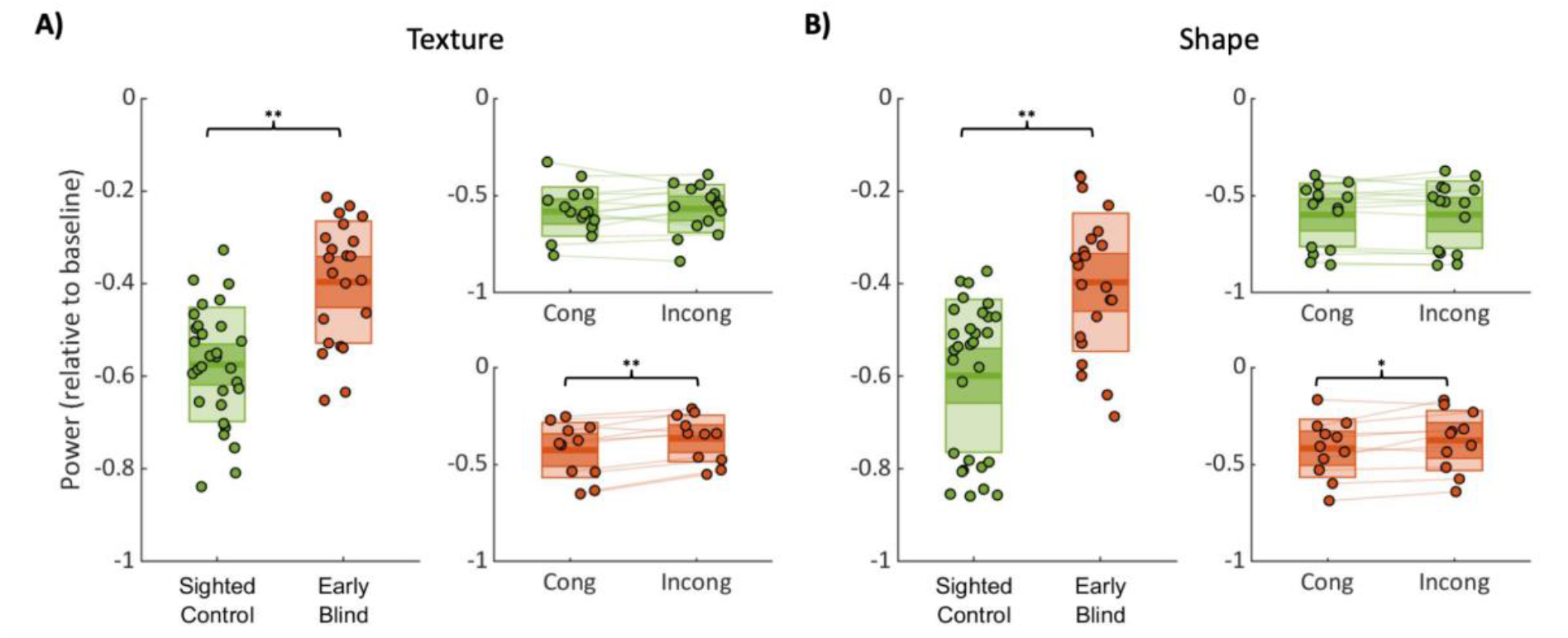
Power values for the frequencies and times of interest at the middle occipital cortex for the texture (A) and shape (B) discrimination tasks. The left graph shows the (baseline-corrected) power for all the trials (congruent and incongruent) separately for the Sighted Control (green) and Early Blind (red) groups. The right-top graph shows the (baseline-corrected) power for the Sighted Control group, separately for congruent and incongruent conditions. The right-bottom graph shows the same information for the Early Blind group. Notice that alpha activity is modulated by the congruency/expectancy of the trial in the Early Blind (but not in the sighted) group in both tactile tasks. *p < 0.005, **p < 0.01.

Similarly, when evaluating the shape discrimination task the ANOVA showed significant effects of group (F_1,24_ = 10.1, p = 0.004), condition (F_1,24_ = 5.8, p = 0.024), and a significant interaction between them (F_1,24_ = 5.1, p = 0.0332). The effect on condition appeared again in the Early Blind individuals only (p = 0.0303), where a larger desynchronization was observed in the Congruent condition. We did not find any differences between conditions in the Sighted Control individuals. These results are shown in Figure 4B.

## DISCUSSION

The present study investigated whether posterior alpha oscillations and their role in gating attention depend on visual experience by assessing whether different attentional conditions could modulate alpha activity in visually deprived people in two (texture and shape discrimination) haptic tasks. Confirming previous observations, alpha rhythms in parieto-occipital areas were significantly reduced in EB individuals in both haptic tasks. Notably, the source reconstruction analysis revealed that the origin of group differences was in the middle occipital lobe. Interestingly, alpha oscillatory activity in that region was modulated by the expectancy of the condition only in the EB group, with congruent trials evoking higher alpha desynchronization than the incongruent condition.

Our observation that alpha oscillatory activity in the middle occipital lobe was modulated by the expectancy of the trial only in the EB group -where a higher alpha suppression was evoked by congruent trials compared to incongruent trials in both discrimination tasks-suggests that the role of alpha oscillations in gating attention does not depend on visual experience. Posterior alpha band activity has been systematically associated with visual attention in sighted people. For instance, parieto-occipital alpha oscillations have been proposed to reflect anticipatory states of visual attention (Foxe et al., 1998; Rihs et al., 2009) and distractor suppression (Kelly et al., 2006; Rihs et al., 2007, 2009). Particularly, alpha-synchronization is thought to be a key component of selective attention, serving active suppression of unattended positions during visual spatial orienting (Rihs et al., 2007). In addition, the direction of attention (predicting spatial biases in imminent visual processing) can be estimated from alpha band lateralization (Kizuk & Mathewson, 2017; Rihs et al., 2007; Thut, 2006) and alpha power reductions have been associated with improved performance (Min et al., 2008; Rohenkohl & Nobre, 2011). In line with the proposed role of alpha oscillations in sighted people, our results show for the first time that alpha oscillations are modulated by attentional demands in blind population and they may play an important role in tactile attention. Therefore, we hypothesize that alpha oscillations maintain the function proposed for visual attention in sighted population but switch to the tactile domain in case of early blindness (Dormal & Collignon, 2011). Our results relates to previous event-related potential studies suggesting that the occipital cortex may be involved in attentional processing and the detection of auditory changes in blindness (Kujala et al., 2005; Liotti et al., 1998). This idea is reminiscent of the observation that occipital regions maintain their computational specialization following the loss of vision, but shift their preferred input modality (e.g., from vision to touch), taking advantage of the underlying computational specialization of that area. For instance, the human visual motion-selective area hMT+ has been found to respond to tactile (Beauchamp et al., 2007; Blake et al., 2004; Jiang et al., 2015; van Kemenade et al., 2014) and auditory motion (Battal et al., 2022; Dormal et al., 2016; Poirier et al., 2006; Wolbers et al., 2011) in case of early blindness. Similarly, ventral occipital regions typically showing category-selective responses to some domain of vision in the sighted were found respond to analogous categories in non-visual modalities in congenitally blind people (Mattioni et al., 2020; Pietrini et al., 2004; Striem-Amit & Amedi, 2014).

The suggestion that alpha activity may be involved in tactile processing in case of early blindness, contrasts with previous suggestions on the role that alpha activity plays in this population. Alpha reductions -compared to sighted controls-have been systematically observed while EB individuals were at rest (Adrian & Matthews, 1934; Hawellek et al., 2013; Noebels et al., 1978; Novikova, 1973), sleeping (Bertolo et al., 2003), during haptic tasks (Kriegseis et al., 2006; Schubert et al., 2015) or during mental imagery tasks (Kriegseis et al., 2006). Consequently, posterior alpha oscillations were considered to depend on structured visual input (Bottari et al., 2016; Kriegseis et al., 2006; Novikova, 1973) and even minimal visual experience (e.g., congenital but incomplete cataracts) was shown to set up the neural mechanisms for alpha generation, whereas the lack of such inputs immediately after birth seemed to result in permanent impairments (Bottari et al., 2016; Innocenti et al., 1985; Novikova, 1973). These reductions were proposed to reflect a decrease of inhibitory processes as a result of reduced thalamic connectivity (Shimony et al., 2006; Singer & Tretter, 1976) and the reduced activity of pyramidal cells, which are modulated by the inhibition of fast-spiking inhibitory interneurons (Buffalo et al., 2011; Jensen et al., 2012). Importantly, the variety of the tasks showing reduced alpha activity in visually deprived population led to suggest that alpha rhythms were not task-specific and did not depend on ongoing sensory or cognitive processes (Kriegseis et al., 2006). Even if the overall reduction of alpha activity that we observe in the EB group fits well with this previous research, our results showing that tactile attention can modulate the posterior alpha band activity challenge the idea that alpha rhythms are not sensitive to task-dependent factors.

Moreover, the observed increased alpha desynchronization in the congruent condition is consistent with the notion that the iterative checking of the expected features requires higher attentional resources than the detection of a mismatch, and that such resources may be allocated via the engagement of task-relevant regions, in this case, the middle occipital cortex of early blind individuals for processing haptic information. The recruitment of middle occipital areas for the processing of tactile attention is not surprising, since the involvement of posterior parieto-occipital regions in blind people during tactile processing -due to crossmodal plasticity-has been previously described (Amedi et al., 2003; Jiang et al., 2015; Sadato et al., 1998, 2004; van Kemenade et al., 2014). Interestingly, the recruitment of posterior brain regions is usually reflected in a relatively higher metabolic activity or blood flow in the occipital cortex, which in turn is negatively correlated with alpha power (Sadato et al., 1998). Regarding the higher desynchronization in the congruent condition, we hypothesize that the iterative checking of the expected features elicits a higher alpha suppression as it requires additional attentional resources compared to the incongruent condition (where a single unexpected feature is enough to discard the expected object). This idea is in line with studies supporting that the function of alpha suppressions is to enhance the neural signal-to-noise ratio (Hanslmayr et al., 2016; Mitchell et al., 2009) and previous work showing that alpha power decreased when stimulus-specific information increased (Griffiths et al., 2019). Likewise, additional resources have been associated with greater alpha band reductions when increasing task difficulty (Pfurtscheller & Lopes da Silva, 1999) or in in older (compared to young) participants during haptic object recognition tasks, suggesting a higher cognitive effort needed to perform the task (Pineda, 2005; Sebastián et al., 2011).

Altogether, our results bring a new understanding to the role that alpha oscillatory activity plays describing plasticity mechanisms in blindness, indicating that alpha activity may reflect tactile attention in blind individuals through the engagement of task-relevant regions such as the middle occipital cortex.

## ACKNOWLEDGEMENTS

The authors thank David Cucurell for his help during the recordings. This work was supported by a Spanish government grant to ARF (PSI2015-69178-P), a predoctoral grant (MINECO-FPI program) from the Spanish government awarded to AGA, project IJC2020-042904-I,MCIN/AEI/10.13039/501100011033 from the Spanish government awarded to AGA, by the Basque Government through the BERC 2022-2025 program, by the Spanish State Research Agency through BCBL Severo Ochoa excellence accreditation CEX2020-001010-S, by the Belgian Excellence of Science (EOS) program (Project No. 30991544) and by Flagship ERA-NET grant SoundSight (FRS-FNRS PINTMULTI R.8008.19).

## Notes

**COMPETING INTERESTS** The authors declare no disclosure of financial interests and potential conflict of interest.

### Competing Interest Statement

The authors have declared no competing interest.

